# Rapid Evolution of Plastic-degrading Enzymes Prevalent in the Global Ocean

**DOI:** 10.1101/2020.09.07.285692

**Authors:** Intikhab Alam, Nojood Aalismail, Cecilia Martin, Allan Kamau, Francisco J. Guzmán-Vega, Tahira Jamil, Afaque A. Momin, Silvia G. Acinas, Josep M. Gasol, Stefan T. Arold, Takashi Gojobori, Susana Agusti, Carlos M. Duarte

## Abstract

Estimates of marine plastic stocks, a major threat to marine life (1), are far lower than expected from exponentially-increasing litter inputs, suggesting important loss factors (2, 3). These may involve microbial degradation, as the plastic-degrading polyethylene terephthalate enzyme (PETase) has been reported in marine microbial communities (4). An assessment of 416 metagenomes of planktonic communities across the global ocean identifies 68 oceanic PETase variants (oPETase) that evolved from ancestral enzymes degrading polycyclic aromatic hydrocarbons. Twenty oPETases show predicted efficiencies comparable to those of laboratory-optimized PETases, suggesting strong selective pressures directing the evolution of these enzymes. We found oPETases in 90.1% of samples across all oceans and depths, particularly abundant at 1,000 m depth, with a strong dominance of *Pseudomonadales* containing putative highly-efficient oPETase variants in the dark ocean. Enzymatic degradation may be removing plastic from the marine environment while providing a carbon source for bathypelagic microbial communities.

---

Exponential growth in plastic production along with poor waste management practices have led to over 150 Tg plastic waste delivered to the ocean since 1950 (1), where it harms marine life, from zooplankton to whales (5). Synthetic plastic polymers are derived from oil hydrocarbons and designed to be durable in the environment (6, 7), being largely resistant to microbial degradation. However, a newly-evolved plastic-degrading enzyme, polyethylene terephthalate hydrolase (PETase), was recently discovered in a Japanese waste processing plant (8). This PETase was inferred to have evolved since 1970, when the polyethylene terephthalate polymer was introduced at industrial scale (9). The PETase catalyzes the hydrolysis of polyethylene terephthalate (PET) plastic to monomeric mono-2-hydroxyethyl terephthalate (MHET), which is ultimately broken down into non-hazardous monomers, terephthalate and ethylene glycol, by MHETases. The efficiency of currently known PETases seems, however, to be low, and it has been suggested that in the environment it only achieves a breakdown from micro-to nanoplastics, rather than full degradation (10).

The discovery, and subsequent biotechnological improvement (11) of this spontaneously-evolved plastic-degrading enzyme offers hope that plastic waste can be degraded in waste facilities, thereby preventing it from entering the ocean (12). However, this still leaves the fate of the > 150 Tg plastic waste already delivered to the ocean unresolved. Recent estimates show that 99% of the plastic that entered the oceans cannot be accounted for, since inventories of plastic floating on the surface of the ocean account to ~ 1% of the expected load (2), pointing at the operation of processes responsible for major losses of plastic at sea (3). One such process may involve the microbial degradation of plastics. Microplastics, the numerically dominant marine debris, have developed a characteristic microbial community, referred to as the marine plastisphere (10). If the PETase enzyme evolved in waste treatment plants from ancestral *Ideonella sakaiensis* (*I. sakaiensis*) genes (8), and provided the current prevalence of polyethylene terephthalate as debris in the marine environment, it is possible that other PETases may have evolved independently in plastisphere ocean microbes as well. The huge population of prokaryotes in the global ocean, estimated to contribute 10^29^ cells distributed among 10^10^ different OTUs (13), provides enough population size, genetic diversity and rate of growth for mutations(14) to independently evolve PETase activity, which would enable microorganisms to access a novel, widely available organic substrate. Indeed, a recent assessment of PETase abundance in marine and terrestrial environments, confirmed the presence of the PETase gene in 31 of the 108 environmental marine samples examined, all from surface waters (4). However, a global assessment of the presence, abundance and diversity of PETases in the whole ocean, extending to the deep ocean, the largest microbial habitat in the biosphere (15), is yet to be conducted.

## Results

Here we report the prevalence of PETases in the global ocean microbiome and provide a phylogeny suggesting Oceanospirillales to be the parental clade leading to 68 identified variants of marine PETase. We do so on the basis of a thorough exploration of PETase through gene catalogs from two global-ocean expeditions, *Tara* Oceans (16) and Malaspina (17), delivering 416 metagenomes collected from 204 individual ocean locations across the Indian, Pacific and Atlantic Oceans from surface to the deep sea (Fig. 1).

**Fig. 1.**
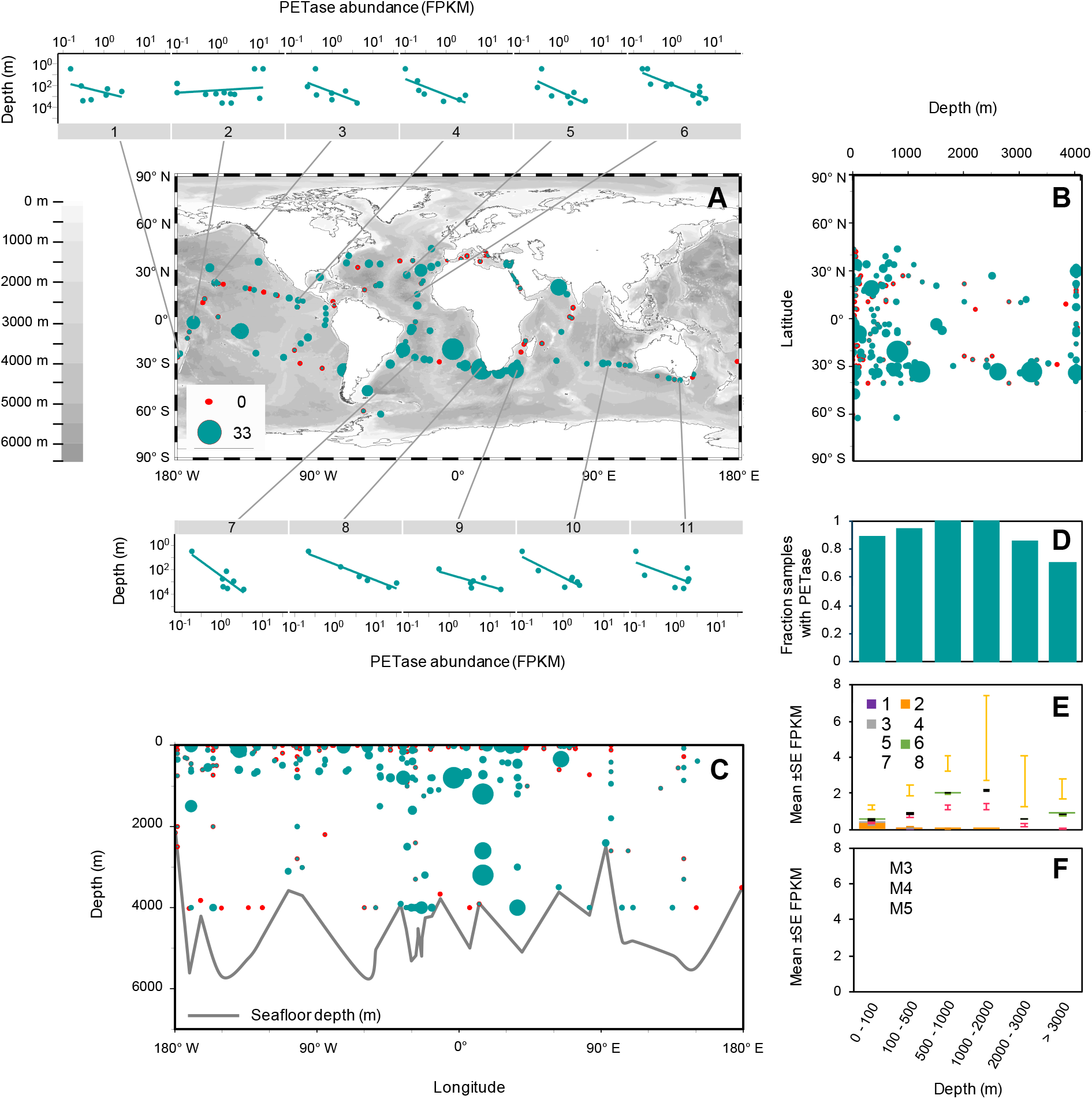
(**A)** Global distribution of PETase (in FPKM) in marine metagenomes. The scale also applies to panels B and C. The 11 small plots show the distribution of PETase genes across depth (m) in 11 profiles collected during the Malaspina Expedition. **(B)** Distribution of PETase along depth and latitude and **(C)** along depth and longitude. The grey line in panel C shows the seafloor depth (m) as it compares to the deepest sampling point. **(D)** Fraction of samples with PETase and mean ±SE PETase abundance (in FPKM) per **(E)** taxonomic group and **(F)** Abundance of oPETases assigned with Motif groups M3-5 (set of motifs designed to capture PETase characteristic catalytic triad and other features) at 6 depth layers (0 - 100 m, 100 - 500 m, 500 - 1000 m, 1000 - 2000 m, 2000 - 3000 m and > 3000 m). Taxonomic groups, identified as numbers from 1 to 8 in panel E, correspond to the following bacterial orders: 1. Candidatus Poribacteria_UO; 2. Flavobacteriales; 3. Gammaproteobacteria_UO; 4. Gemmatimonadetes_UO; 5. Oceanospirillales; 6. Phycisphaerales; 7. Planctomycetales; 8. Pseudomonadales.

Using the KAUST Metagenomic Analysis Platform (KMAP) we queried our pre-annotated *Tara* Oceans and Malaspina gene catalogs for the presence of *Ideonella sakaiensis* PETase (EC 3.1.1.101)-like enzymes containing the Dienelactone hydrolase (DLH) domain. DLH is a generic α/β-hydrolase domain present in many hydrolases, but in the case of PETase it contains a signature catalytic triad (https://pfam.xfam.org/protein/PETH_IDESA, Fig 2a) (4). The DLH domain is present in more than 17,000 sequences in the Pfam database (https://pfam.xfam.org/family/PF01738, accessed Feb 2, 2020). However, most of these sequences are related to genes coding for enzymes involved in the degradation of hydrocarbons. Only one DLH form, harboring specific substitutions in the catalytic triad (motif residues S, D and H) has been experimentally confirmed to degrade PET (4), Fig 2a).

**Fig. 2.**
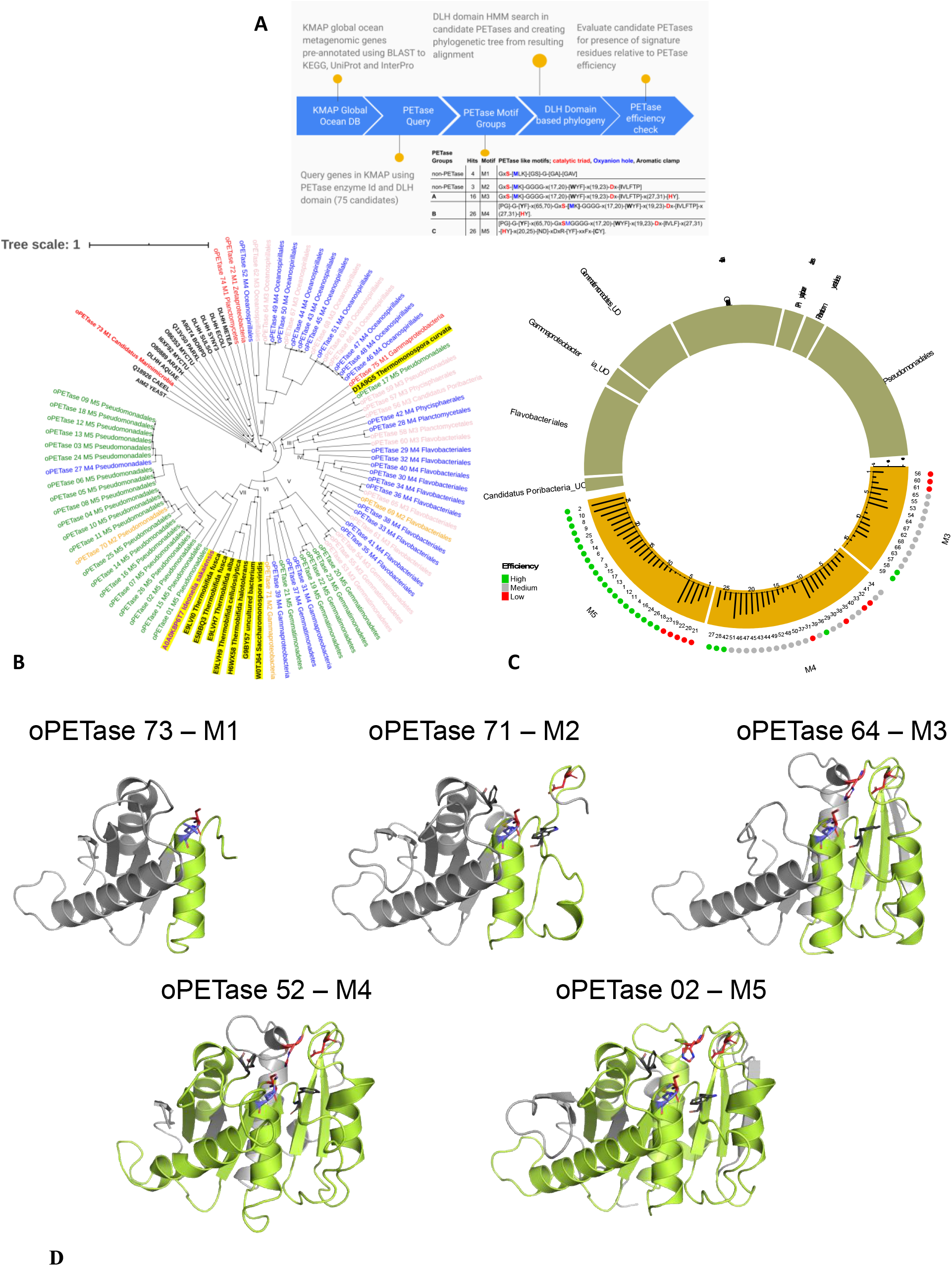
Overview, Phylogenetics and efficiency analysis of candidate Ocean PETases (oPETases). **(A)** An overview of our workflow in exploring oPETases. **(B)** Phylogenetic analysis of DLH-containing PETases. Here sequence names are colored according to associated Motifs M1-5 (M1 in red, M2 in orange, M3 in pink, M4 in blue and M5 in green). DLH seed sequences are in black, similarly known PETases are in black but highlighted in yellow, the original *I. sakaiensis* PETase sequence is in purple and highlighted in yellow. Roman numbers indicate the 7 clades in which PETases are grouped **(C)** Analysis of candidate oPETases is shown for motif groups M3, M4 and M5 (lower half of the circle) presenting potential for efficiency (outer lower circle) as green, grey and red dots depicting high, medium and low efficiency categories, respectively. For potential efficiency to degrade PET, based on the evaluation of only critical residues, see Suppl. Table S2 and efficiency scoring scheme. Total gene abundance for each of these oPETases in ocean is shown in respective bar graphs with orange background. Associated taxonomic information (upper half of the circle) is linked to oPETase motif groups through colored ribbons (M3 is green, M4 is blue and M5 is shown in pink). (D) An illustration of protein structures representative of each motif category, with the corresponding motif M1 to M5 colored in lime green. The models were obtained by SWISS-MODEL based on the crystal structure of IsPETase in complex with HEMT (PDB ID 5xh3) as a template. Key residues are shown as stick models, color-coded as follows: red, the catalytic triad; blue, the Met residue involved in the oxyanion hole; dark grey: residues forming the aromatic clamp, where [YF]87 is part of both the aromatic clamp and the oxyanion hole.

**Fig. 3.**
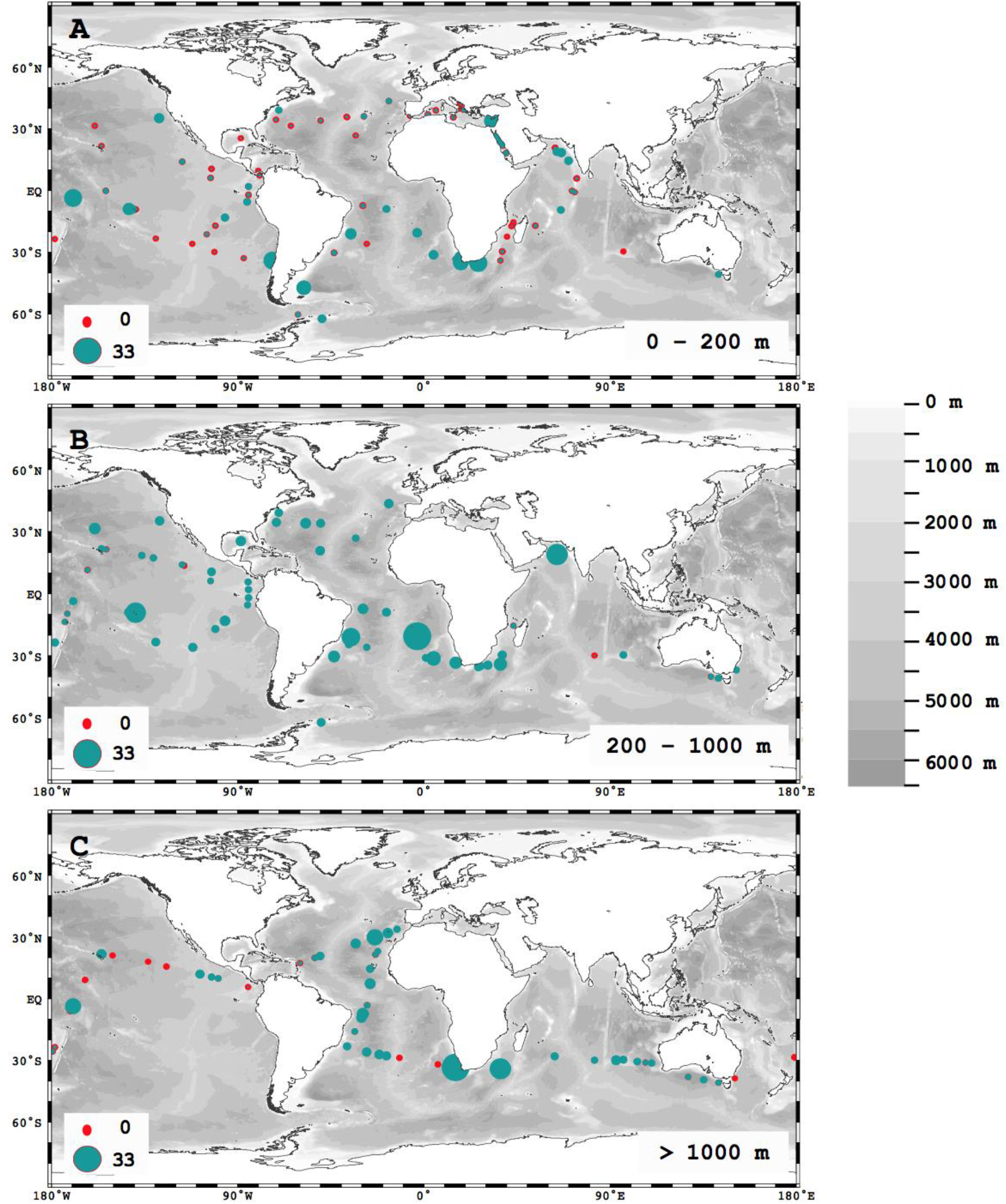
Global distribution of PETase (in FPKM) in marine metagenomes of **(A)** surface (0 - 100 m), **(B)** mesopelagic (200 - 1,000) and **(C)** deep waters (> 1000m).

### Diversity and efficiency of oceanic PETase

Our analysis yielded 75 candidate PETases in the *Tara* Oceans and Malaspina gene catalogues. These gene catalogs were annotated based on full sequence comparisons to public databases. Based on manual inspection of alignments from these 75 oceanic candidate oPETases and 8 known homologs (4) of *I. sakaiensis* PETase we defined five overlapping generic to specific PHI-BLAST searchable motifs, M1-5 (Fig 2a, 2d). Motifs M1 and M2 detect PETase-like sequences but miss the catalytic triad, the main component of known PETases (4). Motif M3 include the catalytic triad, conserved central motif GGGG as well as the oxyanion hole residues and the aromatic clamp. Two exceptions are oPETase 72 and 75 where catalytic triad appears to be available but the conserved motif GGGG is modified to GG-[AG]-[GA]. M4 is similar to M3 but contains the [PG]-G-[YF] motif that defines an additional aromatic clamp and oxyanion hole upstream of the catalytic triad. M5 is the largest motif. It includes M3 and M4 and additionally contains a conserved DxDxR(Y)xxF(L)C sequence, preceding a region providing thermostability to the enzyme (Fig 2a, 2d). Because of the requirement for the catalytic triad of PETases degrade PET, putative PETases containing only motifs M1 and M2 were discarded. Only those with motifs including the catalytic triad (M3, M4 or M5) were considered as functional PETases. This selection yielded 68 variants of putative functional PETases present in the *Tara* Ocean and Malaspina gene catalogues (Suppl. Table 1). Alignments of these oPETases from motif groups M3-5 and shown in Figure 2-suppliment 1.

Scoring of candidate oPETases (see Methods and Suppl. Table S2-3) consistently designated most of the oPETases containing Motif 5 as having a predicted efficiency score close to or slightly above that of the reference *I. sakaiensis* PETase (4), which also contains M5 (Suppl. Table S2). Most of the key residues that are essential for PETase activity are largely conserved across the 68 functional variants. Most of the oPETase variants predicted to be efficient contained M5 (Fig. 2c). Moreover, we found that 20 of the newly identified oPETases, many of which had the M5 motif, already displayed residue substitutions that were identified as enhancing PETase activity in engineered laboratory variants of *I. sakaiensis* PETases (18) (Suppl. Table S3).

To support and inform the scoring process, high-confidence structural homology models were alos produced for all putative PETases, using SWISS-MODEL (19), based on up to 50% sequence identity with the crystal structure of the IsPETase in a complex with HEMT (PDB ID 5xh3 (20)). The structural analysis of potential oPETases can be described using one of the high scoring oPETase example such as oPETase_02. This selected putative PETase contains the catalytic triad (S160, D206, H237) and the disulfide bond linking residues C203 and C239, demonstrated to be essential for PETase function (20, 21)(Fig. 2d). In addition, it harbors two modifications that were shown to increase the catalytic activity of IsPETase (22), namely R90A and L117F. The R90A modification is thought to reduce the steric hindrance around the active site and increase the hydrophobic area to facilitate the accumulation of substrate near the substrate-binding cleft. The L117F modification stabilizes Y87 (F87 in this case), which enhances the interaction between the enzyme and the substrate, as shown by a lower Michaelis constant, *Km* of the L117F mutant as compared to the wild-type PETase. The S214H variant found in some oPETases has been seen to reduce PETase activity, possibly because the larger His restricts the movement of the adjacent W185 (20). Also, other variants that have not been experimentally tested were found in these oPETases (see Supp. Table S1), and their effect on the catalytic activity of the protein remains uncertain. The variants Y87F, T88V and S238Y/F increase the hydrophobicity around the active site, which is expected to be beneficial for PETase activity, as were other substitutions increasing the hydrophobicity of this region, as they may facilitate the interaction of the enzyme with the substrate. The R280Q substitution in most of the higher-scoring variants is conservative and not expected to influence catalysis, and other important positions such as W159, W185 and M161 remain unchanged in these variants. Overall, the compound effects of these modifications make these variants good PETase candidates with a chance of having a PET degrading activity close to or potentially better than IsPETase. It is however important to note that our predicted efficiency scores of these oPETases to rank potentially better or worse cases are adapted based on the presence of favorable or inhibiting mutations only, and that an experimental validation is needed to confirm real efficiency of these oPETases.

### Phylogeny and evolution of efficient oceanic PETase

We established a phylogenetic tree of all PETase sequences, including all Motifs 1 to 5, to investigate the evolutionary pathway from the ancestral DLH domain to fully functional PETase domains including the signature catalytic triad. The phylogenetic tree included the 75 candidate PETases identified in the *Tara* Oceans and Malaspina gene catalogues, the original *I. sakaiensis* PETase sequence (8), the eight land-derived homologues reported from the uniprot database (4), and 12 non-PETase DLH domain core sequences from the Pfam database as an outgroup (Fig. 2b). The resulting phylogenetic tree identified the 12 non-PETase DLH domain core sequences together with two out of three of those containing Motif 1 as a parental group from which PETases evolved, hereafter defined as clade I (Fig. 2b). All other clusters defining six additional taxonomic PETase clades of fully-functional PETases containing M3 to M5 (Fig. 2b), as well as three sequences containing non-functional PETase M2 motifs, interspaced among M3 to M5-containing sequences (Fig. 2b). Clade II is dominated by Oceanospirialles, Clade III contains a mixture of taxa, Clade IV contains Flavobacteriales, Clade V contains Gemmatimonadetes and Gammaproteobacteria, Clade VI contains the eight previously defined homologues of *I sakaiensis* PETase from land microbes (4), with an additional such homologue (thermomonospora curvata, D1A9G5) clustering with clade III (Fig. 2b). Cluster VII contains the original *I. sakaiensis* PETase sequence along with Pseudomonadales sequences, all except two, containing the Motif 5 (Fig. 1b).

In summary, the order of clusters in the phylogenetic tree represents a putative evolutionary progression from DLH-containing ancestors to the eight bacterial orders containing putatively functional oPETases (Fig. 2b). Specifically, the phylogenetic tree obtained suggests that PETases evolved from M1 and M2 DLH-containing sequences of genes involved in the degradation of naturally-occurring hydrocarbons present in the ocean prior to synthetic PET addition to the ocean. The PETases appear to have evolved in parallel in six major independent clades (clades II to VII), all of which include functional PETase motifs (M3, M4 and M5). The phylogenetic tree shows an enrichment of the clades in Motif 5 from clade V, where it first appears, to the clade VII, where all (except one *Pseudomonadales* sequence) include M5. Our analysis points at *Pseudomonadales* containing efficient variants of PETase, as the fundamental outcome of the processes driving the evolution of efficient PETase in the ocean, here oPETase variants 2 and 10 were found to be most abundant (Fig 2c).

### Distribution and abundance of PETases in the global ocean

PETases were prevalent in marine metagenomes, present in 90.1% of the samples assessed, across all oceans and depths sampled (Fig. 1). PETases were present from the surface to the deep sea (Fig. 1b, c), with both prevalence and abundance reaching a maximum at the base of the permanent thermocline (1,000 m depth, Fig. 1d, e), thereby showing a tendency to increase toward the ocean interior (mixed-effect model, slope for depth = 2.979, P = 0.0133, 11 plots in Fig. 1a). This analysis is based on 11 sampling points, where samples were collected at different depths leading to 11 profiles, 10 of which show that PETase abundance increases with depths. PETase abundance in the upper ocean (< 200 m) was highest around South America, South Africa, India, French Polynesia and the Red Sea (Fig. S2a), and PETase abundance in the deep-sea was greatest in the Atlantic Ocean, followed by the Indian Ocean (Fig. S2). Some of these areas, including Southern Africa (23) and Indian subcontinent (24), have been reported to be areas of high abundance of microplastics.

The distribution of PETase contained within *Proteobacteria* was relatively uniform across depth layers (Fig. 1e), however the distribution of PETase contained within *Gemmatimonadetes* increases with depth up to 2,000 m and declines slightly in deeper waters. PETases contained within *Bacteroidetes* were restricted to water depths shallower than 1,000 m (Fig. 1e). PETase-containing *Proteobacteria*, the prokaryote class with the highest abundance and prevalence of PETases, was, in turn, dominated by *Pseudomonadales*, containing efficient M5 motifs, and *Oceanospirillales* orders, with an unidentified order within *Gemmatimonadetes* also contributing considerably to PETase abundance in the mesopelagic ocean (Fig. 1e, Table S1). There was no clear difference in PETase abundance and prevalence between particle attached (> 3 μm) and free-living communities (<3 μm, Wilcoxon Rank Sum Test, W = 2892.5, p-value = 0.46). PETase variants containing the efficient M5 module were dominant across all depth layers (Fig. 1f), with PETase variants containing the M3 variant being the least abundant in oceanic microbial communities (Fig. 1f).

The taxonomy of oPETases differs from that of PETase containing bacteria in other environments. A previous search of 8 homologues of *I. sakaiensis* PETases, all containing the M5 motif, in the public NCBI non-redundant protein database retrieved ~500 PETase-like sequences (4), largely corresponding to microbial communities sampled from terrestrial habitats and related to *Actinobacteria*. An updated search in NCBI (accession date, December 2019), resulted in twice the number of sequences (1138 sequences), also mostly corresponding to soil *Actinobacteria* (1040 sequences). However, ~86 hits appear to be related with proteobacteria (Suppl. Table 4). Our analysis of the phylogeny and abundance of oPETases compared with the results from NCBI clearly show that the PETase-containing ocean microbial community differs, and that PETases likely evolved independently on land and in the ocean.

Our sampling of metagenomes does not allow to directly link the PETase containing microbes to plastic particles. However, our samples are likely to all contain plastic particles. Indeed, as the global metagenomes assessed were retrieved by filtering large volumes of seawater onto filters of different sizes (Fig. 4), any plastic particle, whether marine micro- (< 5 mm) or nanoplastics (< 0.1 μm) (25), present in the water would also be retained. Especially, filters would have retained synthetic fibers (about 0.2 to several mm long and less than 50 μm in diameter), which are the most abundant debris in the ocean, present in the water at concentrations of about 100 to 300 fibers per liter (26). A review of the microbial communities found in the oceanic plastisphere shows that *Pseudomonadales* and *Flavobacteriales*, which our analysis showed that may contain the efficient M5 and M4/M3 PETase, respectively (Fig. 2c), have been detected to form biofilms on marine microplastics sampled in the North Atlantic (27) and North Sea (28). In addition*, Gammaproteobacteria*, which our analysis showed contain the M4-containing PETases (Fig. 2c), have been also reported growing in microbial films on marine microplastics from the Baltic Sea (29), the Caribbean Sea (30), at the coastline of Singapore (31), and through the great Pacific garbage patch (32). However, *Oceanospirillales*, which our analysis showed contain M3 and M4 variants of the PETase, have been identified only in marine plastic-associated bacterial communities in the North Atlantic (27).

**Fig. 4.**
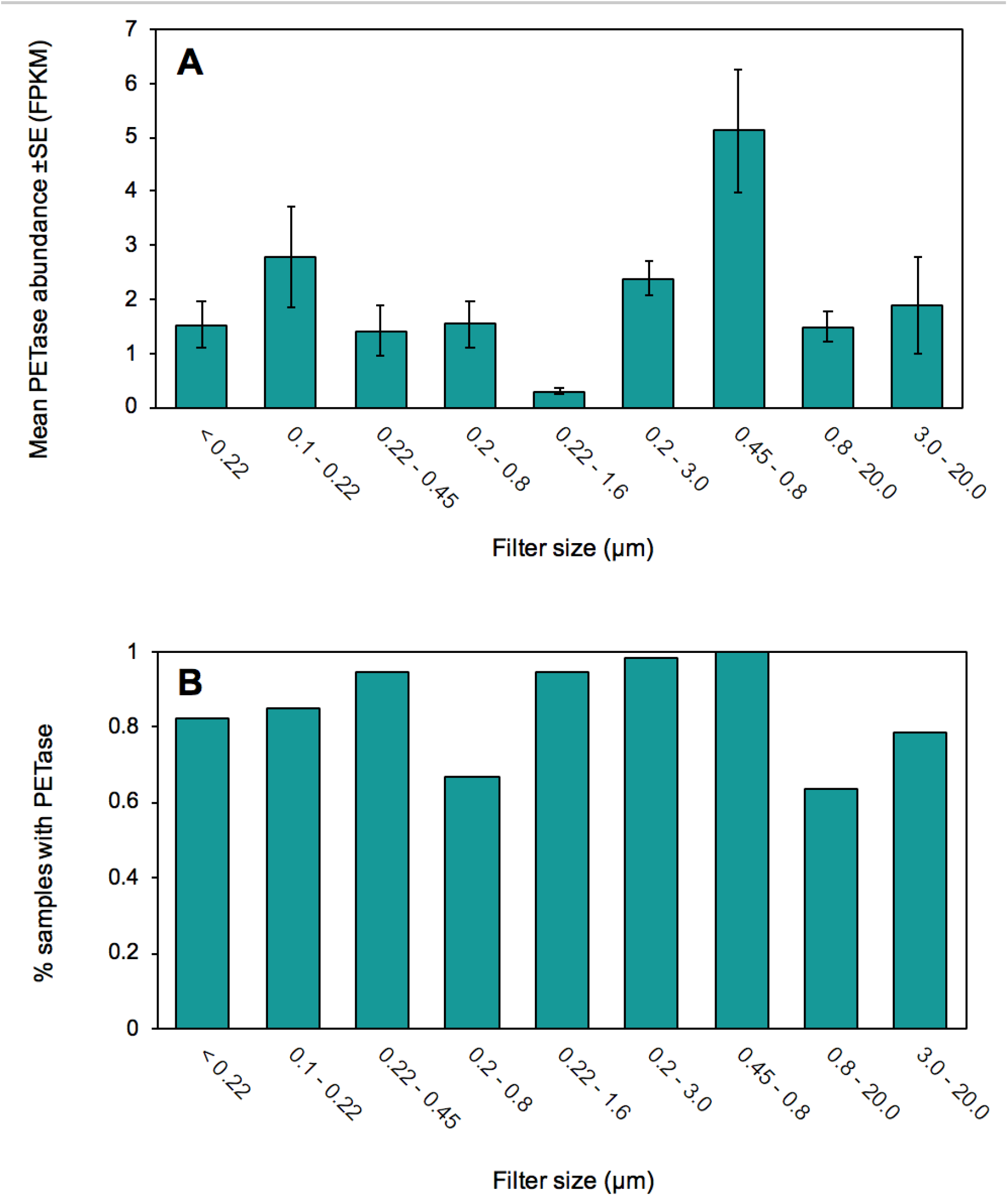
**(A)** Mean PETase abundance ±SE (FPKM) and **(B)** prevalence of samples where PEtase occurred in each of 9 filter sizes (μm).

In summary, the results obtained confirm that PETases are prevalent and abundant in the global ocean microbiome, where they must have evolved recently, following the mass production of polyethylene terephthalate (9). PETases are widespread from the surface to the deep ocean, reaching a maximum abundance and prevalence at about 1,000 m, where *Pseudomonas* has evolved an efficient PETase. PETases were found in high abundance in some areas known to receive high loads of plastic waste, such as waters off the Indian subcontinent, Brazil, and Southern Africa. The phylogeny constructed also shows a putative pathway for the evolution of oPETases from ancestral genes involved in hydrocarbon degradation. The increased efficiency of the PETase along the phylogenetic sequences, along with the high abundance of bacteria containing efficient PETases, points at an ongoing evolutionary process driven by selection pressures providing advantages to bacteria able to use an increasingly available resource, plastic polymers, in the deep ocean, mainly where all other naturally-occurring organic substrates are extremely diluted (33). The widespread prevalence, 90.1%, of PETases across the ocean together with the acute carbon limitation in bathypelagic waters suggests that the ocean microbiome is rapidly evolving to degrade plastic waste, providing a hopeful, nature-based solutions for plastic already in the marine environment.

## Materials and Methods

### Data collection and DMAP based annotation

Metagenome data was retrieved from two global expeditions, the TARA Oceans expedition, which sampled the upper layer of the ocean from sail-boat *Tara* (16) and the Malaspina Expedition, which sampled open ocean waters extending down to 4,000 m depth from spanish R/V *Hesperides* (17). The metagenomic data-sets included 243 metagenome samples from TARA and 173 metagenomes from the MALASPINA expedition, ranging from surface down to 4,000 m depth and including samples from all oceans (Fig. 1). These samples were used to investigate the prevalence of PETase genes. Samples were filtered through different filter sizes (<0.22, 0.1 - 0.22, 0.22 - 0.45, 0.2 - 0.8, 0.22 - 1.6, 0.2 - 3, 0.45 - 0.8, 0.8 - 3.0, 3.0 - 20.0 3 μm) to explore the abundance and prevalence of PETase genes in each fraction and especially in particle attached (> 3 μm) and free-living communities (< 3 μm). Non-redundant Gene catalogs from MP and MD samples were obtained by assembly using MegaHit (34) and gene prediction for each of the samples, followed by pooling of sample specific genes through CD-HIT based clustering (95% nucleotide identity and 80% overlap of the shorter sequence). TARA oceans microbial gene catalog was already available with 40 million genes. MP and MD gene catalogs produced a non-redundant set of 32 and 11 million genes, respectively. These gene catalogs were annotated using DMAP where taxonomic affiliation of every gene is investigated using Least Common Ancestor estimation through BLAST against UniProtKB database. Functional annotation is obtained through BLAST matches to KEGG sequences with defined KEGG Orthologs (KO) and identification of domains using InterPro (35). These Taxonomic and Functional annotations including cross references, such as Enzyme Classification (E.C), Gene Ontology (GO) and BLAST statistics (Evalue, Percent Identity and Percent Coverage) are saved in to Gene Information Table (36) format, ready to be indexed for browsing and comparison through DMAP compare module. To estimate gene abundance counts as Fragments per Kilobase normalized to one Million (FPKM), reads were mapped on to their respective gene catalogs from individual samples in strictly paired mode using bbmap.

### Analysis of plastic degrading-related genes

#### PETase abundance in global ocean metagenomes

We queried TARA, MP and MD gene catalogs to obtain list of non-redundant genes considering enzyme label 3.1.1.101, KEGG Ortholog (KO) K21104 restricted to hits with Pfam signature domain ‘Dienelactone hydrolase family’ with PFAM ID PF01738. Table S1 shows the count of non-redundant genes found as a result of this search. This same query was used in DMAP to obtain gene abundance estimates from individual samples.

#### Phylogeny analysis

A protein sequence set of all non-redundant PETase related 75 genes, obtained through querying DMAP, was combined with *I. sakaiensis* PETase gene, its 8 homologs (from (4)) and as an outgroup 12 seed sequences from DLH domain were included. An alignment of sequences was performed using hmmsearch with DLH domain as a query. Resulting alignment was converted into fasta format and a phylogenetic tree was built using mafft phylogeny option at https://mafft.cbrc.jp/alignment/server/phylogeny.html. The alignment was colored based on different motif groups, *Ideonella sakaiensis* PETase with homologs and seed sequences from DLH domain.

### Efficiency scoring and homology modelling

A scoring system was developed to rank the candidate oPETases, 8 known PET hydrolases, and 12 seed sequences containing the DLH domain. This scoring was made by comparison to the PETase sequence from *I. sakaiensis* (IsPETase), taking into account some of the key residues that are known to contribute to the activity and stability of IsPETase, as well as variants and mutations that have been experimentally tested and yielded either an increase or decrease in its ability to degrade PET (Supplementary Methods and Suppl. Table S2-3). The scoring was then performed with the following considerations: Given the difficulties to predict how the presence of additional varying residues in the sequence with respect to IsPETase might affect the performance of the experimentally tested mutations as reported in the literature, the scoring was applied as a count or sum of the number of positive mutations (i.e. enhancing PETase activity) minus the number of negative mutations (i.e. inhibiting PETase activity). Additionally, a higher weight was given to the residues composing the catalytic triad and the PETase-specific disulfide bond, and a higher penalty to their absence. A labelling of PETases based on resulting score binned into low, medium and high efficiency, depicted by red, grey and green colors, respectively, is shown in Fig. 1c. After several rounds of scoring varying different parameters, such as the number of mutations/key amino acids considered and the weights given to each residue, the ranking of the analyzed sequences was proven to be robust.

## Acknowledgments

We are thankful to KAUST Supercomputing Laboratory (KSL) for providing computational resources need to carry out this work,

## Funding

The research reported in this publication was supported by funding from King Abdullah University of Science and Technology (KAUST), Office of Sponsored Research (OSR), under award number URF/1/1976-21-01, and the Malaspina Circumnavigation Expedition funded by the Spanish Ministry of Economy and Competitiveness (CSD2008-00077),;

## Author contributions

IA and CMD conceived the study, SGA. JMG, TG, CMD and SA contributed data, IA, AK, CMD, NA, CM and TJ analyzed the metagenome data, FJGV, AAM and STA conducted the structural modelling and efficiency assessment, IA, CMD, NA and CM wrote the first draft of the paper and all coauthors contributed to improve the paper and approved the submission;

## Competing interests

Authors declare no competing interests; and

## Data and materials availability

All data are available (see sequence data information in M&M).

---

**Fig 2 Supplementary 1.**
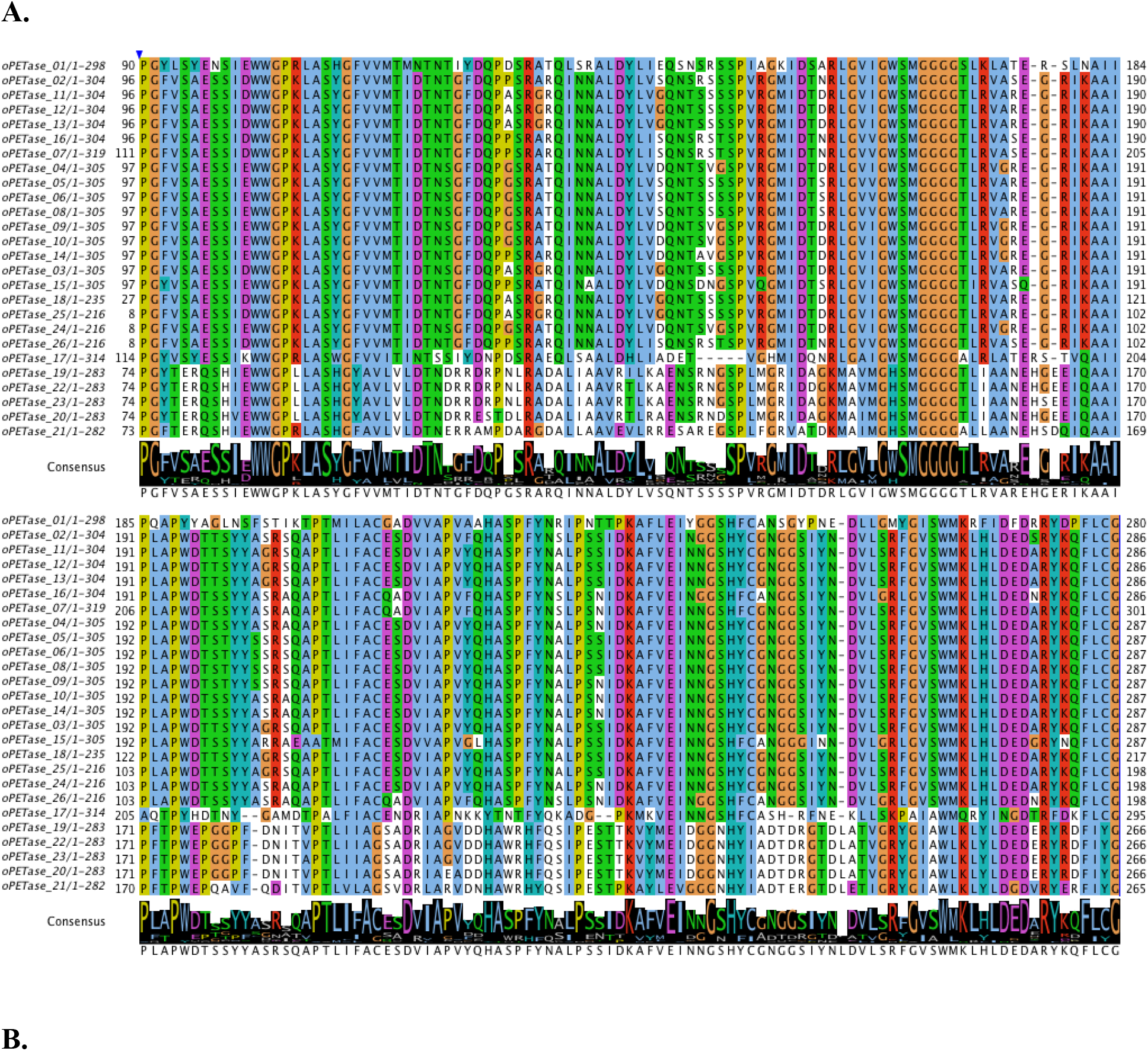
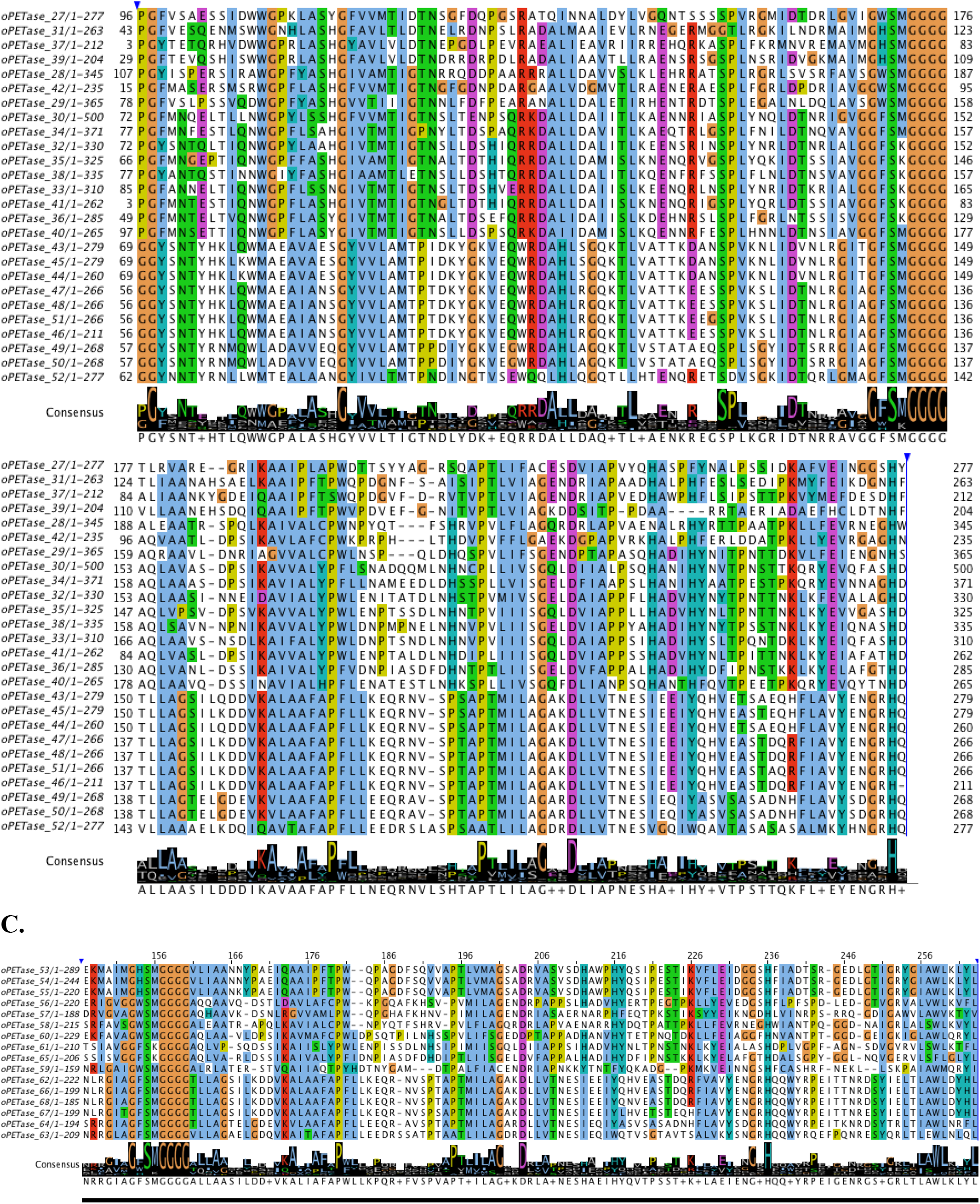
Alignments of ocean PETase groups M5 (A), M4(B) and M3 (C) showing conservation and variation in key PETase residues particularly catalytic tirade, aromatic clamp and oxyanion hole.

